# Effects of protein boosters on antibody responses

**DOI:** 10.1101/2025.02.08.637239

**Authors:** Bakare Awakoaiye, Tanushree Dangi, Pablo Penaloza-MacMaster

## Abstract

SARS-CoV-2 has infected a large fraction of the human population. Currently, most individuals have developed immunity either through vaccination or natural infection. Despite this, SARS-CoV-2 booster immunizations are still recommended to reduce the risk of reinfections, but there is still limited understanding on how different booster vaccine platforms influence antibody responses. We conducted immunological studies in mice to evaluate the boosting effects of different vaccine platforms on antibody responses. C57BL/6 mice were first primed with an adenovirus serotype 5 (Ad5) vaccine expressing the SARS-CoV-2 spike protein. The mice were then boosted with the same Ad5-based vaccine (homologous boosting) or with a protein-based vaccine (heterologous boosting). Interestingly, the heterologous regimen (Ad5 prime + protein boost) elicited superior antibody responses, relative to the homologous regimen (Ad5 prime + Ad5 boost). Similar potentiation of antibody responses was reported when mice were primed with poxvirus or rhabdovirus vectors and then boosted with protein. These findings highlight a potential advantage of protein booster immunizations to potentiate humoral immunity.

## INTRODUCTION

The COVID-19 pandemic has led to widespread global immunity to SARS-CoV-2 through vaccination, natural infection, or a combination of both. However, waning antibody responses over time have been linked to an increased risk of breakthrough infections, underscoring the importance of booster vaccinations to sustain protective immunity ^1^. Despite the broad deployment of booster vaccines, there is still limited understanding of how different vaccine platforms influence antibody responses after boosting.

Adenovirus-based vaccines, including Ad5-based vaccines such as the CanSino and Sputnik V vaccines, and vaccines utilizing other adenovirus serotypes like Ad26 or chimpanzee adenovirus, have been administered to millions of people worldwide and demonstrated efficacy against COVID-19 ^2-4^. Additionally, mRNA vaccines and protein subunit vaccines, which are key components of the global vaccination effort, have played a significant role in curbing the pandemic and are currently authorized for use in many countries, including the United States. In particular, protein vaccines offer the advantage of low reactogenicity compared to other vaccine platforms, making them an appealing option for booster immunizations.

To explore how protein booster vaccines affect immune responses, we conducted studies in mice primed with viral vector vaccines. These mice were subsequently boosted either with homologous viral vector vaccines or with heterologous protein vaccines. Our findings reveal that heterologous boosting with protein vaccines elicits significantly higher antibody responses compared to homologous viral vector boosting. These results provide insights into optimizing vaccine booster strategies to specifically enhance humoral immunity.

## RESULTS

### Priming with Ad5 and boosting with protein elicits superior SARS-CoV-2 specific antibody responses, relative to priming and boosting with Ad5

We first primed C57BL/6 mice intramuscularly with an Ad5 adenovirus vector expressing SARS-CoV-2 spike protein (Ad5-SARS-CoV-2 spike)^5^ or a spike protein vaccine. After approximately four weeks, mice were boosted homologously or heterologously, and immune responses were measured in blood (Fig. 1A). Priming and boosting with the same Ad5 vaccine resulted in a pattern of improved CD8 T cell responses, relative to Ad5 priming and protein boosting (Fig. 1B). However, Ad5 priming and protein boosting resulted in greater antibody responses (Fig. 1C). This enhancement in antibody responses observed in the heterologous prime boost regimen was associated with an increase in plasma cells in the bone marrow (Fig. 2).

**Fig. 1.**
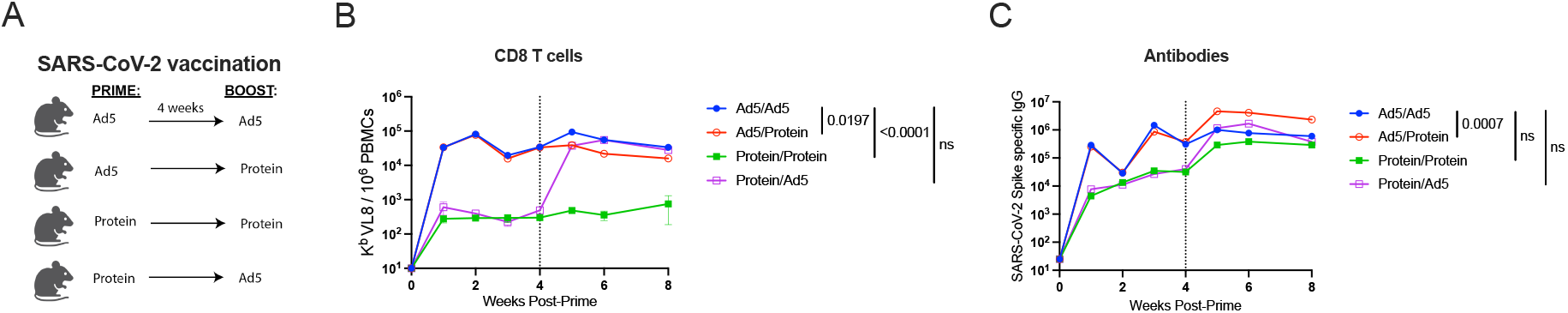
Priming with Ad5-SARS-CoV-2 spike and boosting with SARS-CoV-2 spike protein elicits superior antibody responses, compared to homologous prime-boost vaccine regimen. **(A)** Experimental approach for evaluating how protein boosting affects adaptive immune responses. C57BL/6 mice were primed and boosted with either 10^8^ PFU of an Ad5-SARS-CoV-2 spike vaccine, or 10 µg of spike protein in 1:10 Adju-Phos. **(B)** Summary of SARS-CoV-2-specific CD8^+^ T cell responses (K^b^ VL8+) in PBMCs. Vertical dashed line indicates time of boosting. **(C)** Summary of SARS-CoV-2-specific antibody responses in sera. Data are from five experiments, with n=4-7 mice per group. All data are shown. Indicated *P* values were determined by one-way ANOVA Kruskal-Wallis test with Dunnett’s multiple comparison at the final timepoint. Error bars represent SEM.

**Fig. 2.**
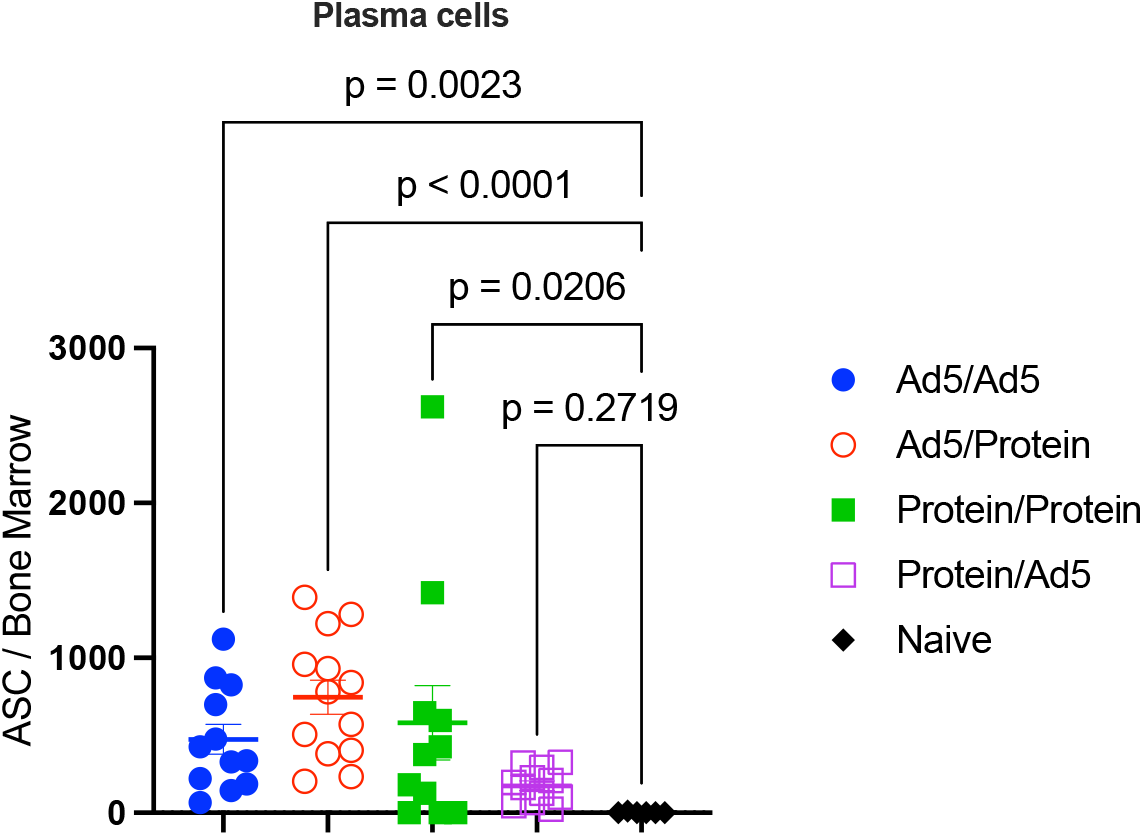
Plasma cells following vaccination. C57BL/6 mice were primed and boosted with either 10^8^ PFU of an Ad5-SARS-CoV-2 spike vaccine, or 10 µg of spike protein in 1:10 Adju-Phos. Antibody secreting cells were enumerated in bone marrow after 20 weeks. Data are from three experiments, with n=3-5 mice per group. Each experiment included 1-2 naïve control mice. All data are shown. Indicated *P* values were determined by one-way ANOVA Kruskal-Wallis test with Dunnett’s multiple comparison. Error bars represent SEM.

### Generalizability to other Ad5 vaccines

The above studies only involved Ad5 and protein vaccines for SARS-CoV-2, so we broadened our analysis to determine whether our findings could generalize to other antigens. We primed C57BL/6 mice intramuscularly with an Ad5 adenovirus vector expressing lymphocytic choriomeningitis virus (LCMV) glycoprotein (Ad5-LCMV GP). After approximately four weeks, mice were boosted homologously with Ad5-LCMV GP or heterologously with the respective protein, LCMV GP, and immune responses were measured in blood (Fig. 3A). Priming and boosting with the same Ad5 vaccine resulted in similar LCMV-specific CD8 T cell responses, relative to Ad5 priming and protein boosting (Fig. 3B). However, the heterologous Ad5 prime and protein boost resulted in significantly greater LCMV-specific antibody responses (Fig. 3C).

**Fig. 3.**
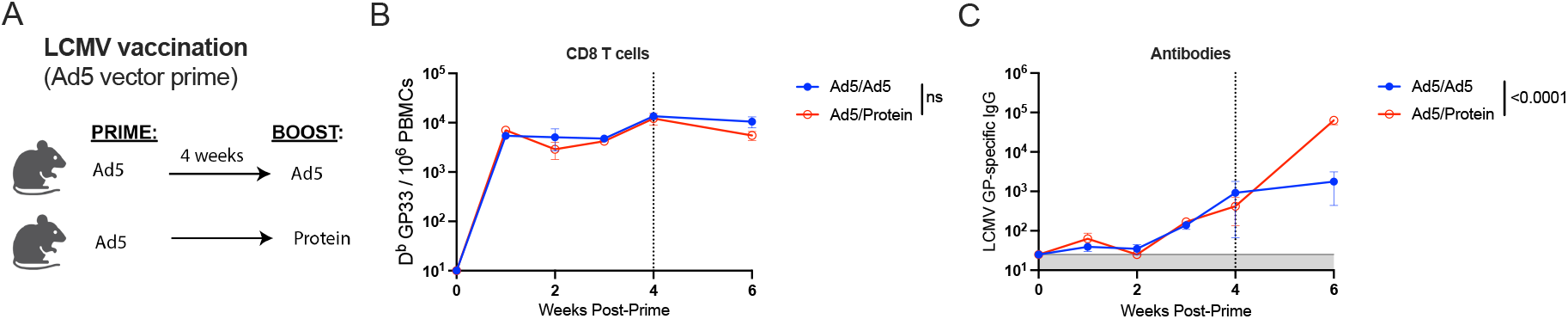
Priming with Ad5-LCMV-GP and boosting with LCMV GP protein elicits superior antibody responses, compared to homologous prime-boost vaccine regimen. **(A)** Experimental approach for evaluating how protein boosting affects adaptive immune responses. C57BL/6 mice were primed and boosted with either 10^8^ PFU of an Ad5-LCMV-GP vaccine, or 10 µg of GP protein in 1:10 Adju-Phos. **(B)** Summary of LCMV-specific CD8^+^ T cell responses (D^b^ GP33+) in PBMCs. **(C)** Summary of LCMV-specific antibody responses in sera. Data are from two experiments, with n=7-8 mice per group. All data are shown. Indicated *P* values were determined by Mann Whitney test. Error bars represent SEM.

### Generalizability to other vaccine platforms

The above studies involved Ad5 and protein vaccines expressing various antigens, and we then explored whether our findings could generalize to different viral vector platforms other than Ad5. We primed C57BL/6 mice intramuscularly with a poxvirus vector (vaccinia virus) expressing Lassa virus glycoprotein (VV-Lassa GP). After four weeks, mice were boosted homologously with VV-Lassa GP or heterologously with the respective protein, Lassa GP, and antibody responses were measured in blood (Fig. 4A). Priming with VV-Lassa GP and boosting with protein resulted in significantly greater antibody responses, relative to the homologous VV-Lassa GP prime boost regimen (Fig. 4B). Similar results were observed with a vaccinia virus vector expressing a human immunodeficiency virus (HIV-1) envelope protein (VV-HIV env) (Fig. 4C-4D), and a modified vaccinia Ankara expressing SARS-CoV-1 spike protein (MVA-SARS-CoV-1 spike) (Fig. 4E-4F).

**Fig. 4.**
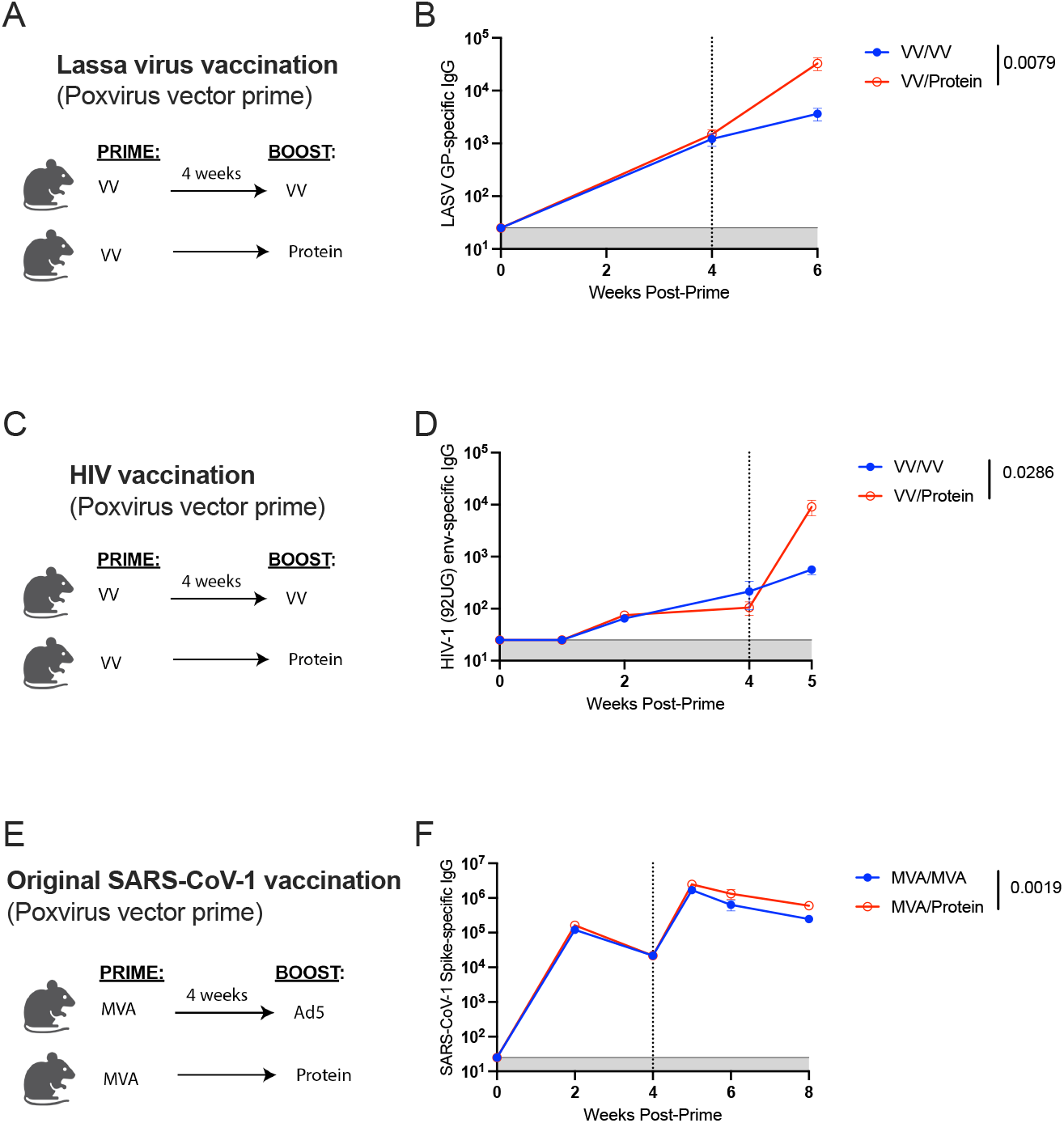
Priming with poxvirus vectors and boosting with protein elicits potent antibody responses. **(A)** Experimental approach. C57BL/6 mice were primed and boosted with either 10^8^ PFU of a VV-Lassa GP vaccine, or 5 µg of Lassa GP protein in 1:10 Adju-Phos. **(B)** Summary of Lassa-specific antibody responses in sera. **(C)** Experimental approach. C57BL/6 mice were primed and boosted with either 7×10^7^ PFU of a VV-HIV Env (strain 92UG) vaccine, or 100 µg of HIV Env protein in 1:10 Adju-Phos. **(D)** Summary of HIV-specific antibody responses in sera. **(E)** Experimental approach. C57BL/6 mice were primed and boosted with either 3×10^6^ PFU of a MVA-SARS-CoV-1 vaccine, or 10 µg of SARS-CoV-1-spike in 1:10 Adju-Phos. **(F)** Summary of SARS-CoV-1-spike-specific antibody responses in sera. Data from B and D are from one experiment, with n=4-5 mice per group; data from F are from two experiments, with n=4-5 mice per group. All data are shown. Indicated *P* values were determined by Mann Whitney test. Error bars represent SEM.

In addition, we investigated whether our findings could extend to a vesicular stomatitis virus vector (VSV) expressing a SARS-CoV-2 spike protein (VSV-SARS-CoV-2 spike). We first primed C57BL/6 mice intramuscularly with VSV-SARS-CoV-2 spike, and after about four weeks, mice were boosted homologously with the same VSV-SARS-CoV-2 spike viral vector vaccine or heterologously with SARS-CoV-2 spike protein, and antibody responses were measured in blood. Priming with the VSV-SARS-CoV-2 spike vaccine and boosting with the spike protein vaccine resulted in significantly greater antibody responses, relative to the homologous prime boost regimen (Fig. 5A-5B). Altogether, these findings suggest that protein boosters are particularly useful to enhance antibody responses.

**Fig. 5.**
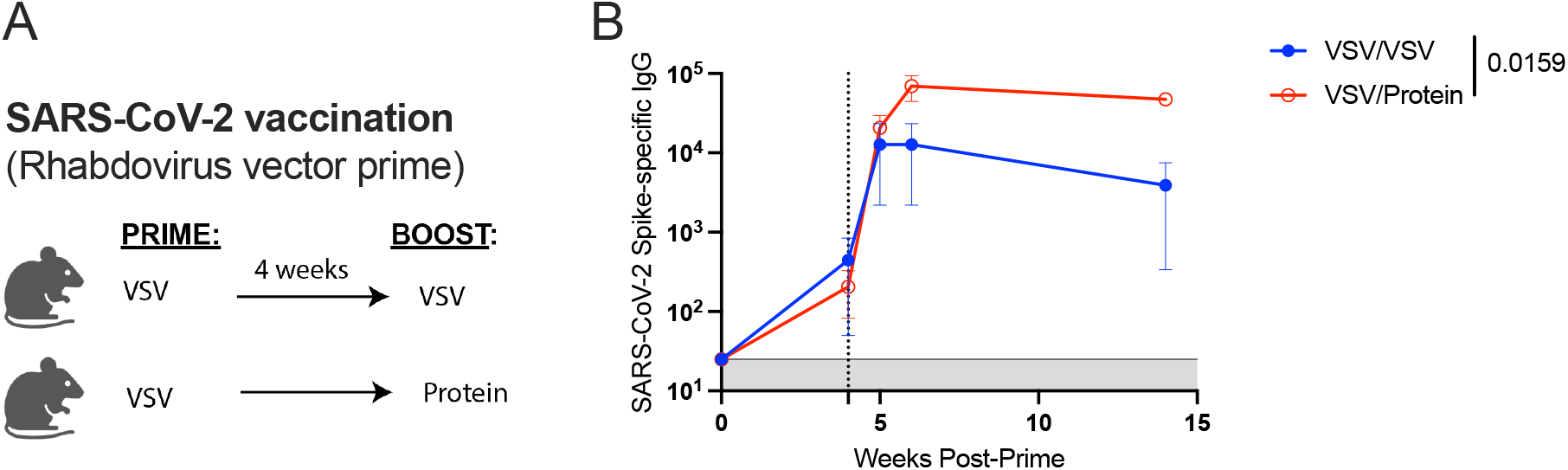
Priming with a rhabdovirus vector and boosting with protein elicits potent antibody responses. **(A)** Experimental approach. C57BL/6 mice were primed and boosted with either 6×10^5^ PFU of a VSV-SARS-CoV-2 spike vaccine, or 10 µg of SARS-CoV-2 spike protein in 1:10 Adju-Phos. **(B)** Summary of SARS-CoV-2 spike-specific antibody responses in sera. Data are from one experiment, with n=5 mice per group. All data are shown. Indicated *P* values were determined by Mann Whitney test. Error bars represent SEM.

## DISCUSSION

Although SARS-CoV-2 vaccines prevent severe disease and death, they do not confer durable sterilizing immunity due in part to viral variants as well as waning antibody titers, warranting the use of boosters. In this report, we show that a heterologous regimen composed of a viral vector prime followed by a protein boost results in superior antibody responses compared to homologous prime-boost with viral vectors. What is the mechanistic basis for this? It is possible that after a viral vector prime, anti-vector immunity could reduce the boosting potential of Ad5. Pre-existing immunity elicited by either active or passive immunization can affect the boosting capacity of vaccines ^6, 7^, and in particular, anti-vector immunity is known to accelerate vaccine clearance and immunogenicity of viral vectors like Ad5 ^8^. It is also possible that soluble protein is more effective at engaging B cell receptors (BCR) on the surface of memory B cells. Another important aspect to highlight in some of our vaccine studies (SARS-CoV-2 vaccination studies), is that the homologous Ad5 prime / Ad5 boost regimen tended to elicit slightly superior CD8 T cell responses than the heterologous Ad5 prime / protein boost regimen. This is likely because viral vectors express their antigens intracellularly, favoring CD8 T cell responses. However, this improvement in CD8 T cell responses with the homologous Ad5 regimen was not statistically significant in the case of Ad5-GP.

Our data with Ad5 vectors expressing different antigens (Ad5-GP and Ad5-HIV) and our data with different viral vector platforms (VV, MVA, VSV) suggest generalizability. We did not conduct SARS-CoV-2 challenge experiments to compare immune protection because in prior mouse studies, vaccinating with the Ad5-SARS-CoV-2 spike vaccine conferred robust protection, making it difficult to detect differences in immune protection between homologous and heterologous Ad5 vaccine regimens. High level of immune protection has also been described with other SARS-CoV-2 vaccines based on adenoviruses ^9-13^. Similarly, the Ad5-LCMV GP vaccine results in very rapid clearance of chronic LCMV Cl-13 ^14^, rendering it difficult to ascertain differences in immune protection between different booster regimens.

However, it is possible that a viral vector prime and protein boost regimen may be useful in individuals who develop suboptimal antibody responses following vaccination, including the elderly. Protein boosters may also be especially effective in the context of HIV vaccination, where antibody responses are believed to be more critical than T cell responses at preventing initial infection and establishment of the viral reservoir. Boosting with protein may also improve uptake and safety in the population, given that protein vaccines induce lower adverse events compared to other vaccine platforms. It is also important to note that while only a small fraction of people in the United States received adenovirus vaccines during the COVID-19 pandemic, these vaccines served as the primary option for many underdeveloped countries. Future studies will determine whether protein boosters have the same effect in the setting of mRNA vaccines. Altogether, these data suggest that protein boosters favor the elicitation of potent antibody responses in hosts that were previously primed with viral vectors, providing insights for rational vaccine design.

## MATERIALS AND METHODS

### Vaccines

We used a non-replicating Ad5 vector that is E1/E3 deleted and expresses SARS-CoV-2 spike protein (strain 2019-nCoV-WIV04) within the putative E1 site ^5, 15^. This vector contains a Cytomegalovirus (CMV) promoter driving the expression of the SARS-CoV-2 spike protein and was used in prior papers ^5, 8^. The other Ad5 vectors have a similar genetic profile, but they express their respective transgene in the putative E1 site, instead of spike. The Ad5 vectors were propagated at the Iowa Vector Core on trans-complementing HEK293 cells (ATCC), purified by cesium chloride density gradient centrifugation, titrated, and then frozen at −80 °C. All the other vectors were propagated in house. The poxvirus vectors were a kind gift from Dr. Bernard Moss (NIH). The rhabdovirus (VSV) vector was a kind gift from Dr. Sean Whelan (Washington University in St. Louis). The Ad5-LCMV GP vector was a gift from Dr. Julie McElrath (Fred Hutchinson Cancer Research Center).

### Mice and vaccinations

6-8-week-old C57BL/6 mice were used. Mice were purchased from Jackson laboratories (approximately half males and half females). Mice were immunized intramuscularly (50 µL per quadriceps) with the respective vaccine diluted in sterile PBS. Mice were housed at the Northwestern University Center for Comparative Medicine (CCM) in downtown Chicago. All mouse experiments were performed with approval of the Northwestern University Institutional Animal Care and Use Committee (IACUC).

### Reagents, flow cytometry, and equipment

Single cell suspensions were obtained from PBMCs and various tissues as described previously ^16^. Dead cells were gated out using Live/Dead fixable dead cell stain (Invitrogen). Biotinylated MHC class I monomers (K^b^ VL8, sequence VNFNFNGL, and D^b^GP33, sequence KAVYNFATC, used and tested in prior studies ^17-22^) were obtained from the NIH tetramer facility at Emory University. Cells were stained with fluorescently labeled antibodies against CD8α (53-6.7 on PerCP-Cy5.5) purchased from BD Pharmingen and CD44 (IM7 on Pacific Blue) purchased from Biolegend. Flow cytometry samples were acquired with a Becton Dickinson Canto II or an LSRII and analyzed using FlowJo (Treestar).

### ELISA and antibody secreting cells assays

Binding antibody titers were measured using ELISA as described previously ^18, 23-25^, but using protein instead of viral lysates. In brief, 96-well flat bottom plates MaxiSorp (Thermo Scientific) were coated with 0.1μg/well of the respective protein, for 48 hr at 4ºC. Plates were washed with PBS + 0.05% Tween-20. Blocking was performed for 4 hr at room temperature with 200 μL of PBS + 0.05% Tween-20 + bovine serum albumin. 6μL of sera were added to 144 μL of blocking solution in first column of plate, 1:3 serial dilutions were performed until row 12 for each sample and plates were incubated for 60 minutes at room temperature. Plates were washed three times followed by addition of goat anti-mouse IgG horseradish peroxidase conjugated (Southern Biotech) diluted in blocking solution (1:1000), at 100 μL/well and incubated for 60 minutes at room temperature. Plates were washed three times and 100 μL /well of Sure Blue substrate (Sera Care) was added for approximately 8 minutes. Reaction was stopped using 100 μL/well of KPL TMB stop solution (Sera Care). Absorbance was measured at 450 nm using a Spectramax Plus 384 (Molecular Devices). SARS-CoV-2 spike protein was made in house using a plasmid that was produced under HHSN272201400008C and obtained through BEI Resources, NIAID, NIH: Vector pCAGGS containing the SARS-Related Coronavirus 2, Wuhan-Hu-1 Spike Glycoprotein Gene (soluble, stabilized), NR-52394. SARS-CoV-1 spike protein was obtained through BEI Resources, NIAID, NIH: SARS-CoV Spike (S) Protein deltaTM, Recombinant from Baculovirus, NR-722. A hybridoma for expressing LCMV GP Cl-13 protein was a kind gift of Drs. Carl Davis and Rafi Ahmed. The same proteins used as vaccines were used as coating antigen for the ELISA. For detecting antibody secreting cells (ASC) we used a protocol from a prior publication ^23^.

### Statistical analysis

Statistical tests used are indicated on each figure legend. In the ELISA data, horizontal shaded lines in figures represent limit of detection. Data were analyzed using Prism version 9 (Graphpad).

